# Liberating links between datasets using lightweight data publishing: an example using plant names and the taxonomic literature

**DOI:** 10.1101/343996

**Authors:** Roderic D. M. Page

## Abstract

Constructing a biodiversity knowledge graph will require making millions of cross links between diversity entities in different datasets. Researchers trying to bootstrap the growth of the biodiversity knowledge graph by constructing databases of links between these entities lack obvious ways to publish these sets of links. One appealing and lightweight approach is to create a “datasette”, a database that is wrapped together with a simple web server that enables users to query the data. Datasettes can be packaged into Docker containers and hosted online with minimal effort. This approach is illustrated using a dataset of links between globally unique identifiers for plant taxonomic names, and identifiers for the taxonomic articles that published those names.

## Introduction

A venerable tradition in taxonomy is compiling and publishing lists of scientific names, whether in printed form or as online databases. If the lists include bibliographic information these lists can also serve as search indices to the taxonomic literature. However, this functionality is often hampered by the use of cryptic citations, and a mismatch between the granularity sought by taxonomists (often page-level) and that used by researchers when citing sources. A practical consequence is that a biologist seeking information about a species may struggle to locate the original taxonomic description, which for many species may be the best (or, indeed, only) source of basic biological data for that species [Page 2016a].

In an ideal world each taxonomic name would be linked to a detailed bibliographic record of where that name was published, and that publication would be available in digital form, as would any subsequent taxonomic revisions [Agosti & Egloff,2009, Page 2016a]. There are notable examples of resources like this for particular taxonomic groups (e.g., World Spider Catalog)[Gloor et al. 2017], but there is no freely accessible resource that covers, for example, all animals, or all plants. There are detailed databases of names that also cite the primary literature, but typically these citations are simply text strings, not actionable digital identifiers.

Motivated by this lack of links I have spent the last few years obsessively collecting digital identifiers for taxonomic publications and linking them to taxonomic names. This project is far from complete, nor is it likely to be in the near future given the continuing discovery of new species, and the increasing number of taxonomic works that are becoming available online. One consequence of this Sisyphean task is that it becomes tempting to simply continue to accumulate links in a local database, forever postponing actually publishing them. This is not a particularly successful career strategy, nor is it helpful to people who might make use of these links. However, publishing sets of links is not necessarily a straightforward task.

One option for publication is to create a custom interface to the links, to make them both discoverable and interesting. Examples include links between the NCBI taxonomy [Federhen 2011] and Wikipedia [Page 2011], or BioNames [Page 2013], which comprises links between animal names and the primary literature. These may be user friendly, but they provide limited functionality, especially if a user wants programmatic access to the underlying data.

Rather than expend effort on developing idiosyncratic solutions, one could simply publish the data to an existing platform. I adopted this approach for the names in the Plant List [http://www.theplantlist.org/], for which I linked a subset of names to publications with Digital Object Identifiers (DOIs) or with a link to JSTOR. This dataset was uploaded to the Global Biodiversity Information Facility (GBIF) [Page 2016c]. GBIF uses the Darwin Core Archive [Wieczorek et al. 2012] as its data format, which does not handle literature particularly well, and literature is not a first class citizen of the GBIF portal. Consequently, uploading this data to GBIF does not make the most of the effort that went into creating the links.

GBIF is a domain-specific data publisher. An alternative may be to publish in a venue with broader scope, such as Wikidata [Vrandečić & Krötzsch 2014, https://www.wikidata.org]. Wikidata is rapidly becoming a useful platform for cross linking scientific data [Burgstaller-Muehlbacher et al. 2016], and has enormous potential. However, because the data is published at the level of individual statements, it becomes difficult to point to who did what in a simple way. In contrast, GBIF treats datasets as both a bundle of data that can be identified as a specific contribution (with a dataset-specific DOI), as well as “unbundling” the data and merging it into a single index.

A particularly appealing route for publishing links would be to treat each link as a “nanopublication” [Groth et al. 2010], which is minimally a single linked data “triple”. Nanopublications have built in mechanisms for provenance and attribution [Kuhn et al. 2017], and have been used to publish large datasets [Queralt-Rosinach et al. 2016]. Because nanopublications are grounded in linked data they will be of most use in communities where linked data has been widely adopted. To date the biodiversity informatics community has shown lukewarm enthusiasm for linked data, despite various calls to exploring its use [Page 2016b], and some working implementations [Michel et al., 2017; Senderov et al. 2018], we have nothing on the scale, say, of Uniprot [The UniProt Consortium 2016]. Hence, despite their attractiveness, publishing the links as nanopublications does not seem to be a way to encourage their reuse, although this may well change in the future.

If we find the three options discussed so far (custom web site, existing data publisher, and nanopublications) unsatisfactory, then it seems that the only remaining approach is simply to deposit the dataset as a “dumb” file in a repository such as Datadryad or Zenodo, minting a DOI to make it citable, and then hope that somebody makes use of it. But multi-megabyte data files are often not the easiest for users to work with, and it might not be obvious to a potential user why the data would be worth them investing time in discovering whether it was useful.

However, other possibilities are emerging. For example, Simon Willison [2017] has recently proposed a lightweight approach to data publishing called “datasettes”. A datasette comprises one or more comma separated value (CSV) files which are merged into a SQLite database. The database and a simple API are bundled together with a web server that can be queried interactively by the user via a web interface. The datasette can be run on the user’s local machine, or easily pushed to a server in the “cloud”, for example using Docker containers. Datasettes and the other approaches listed above are not, of course, mutually exclusive. But the attraction of the datasette is that it makes it easy to publish data that might otherwise either not be published, or might be published as a large “blob” of data whose utility is opaque to its potential users.

In this paper I describe the creation of a datasette for a longstanding but mostly unpublished project on linking plant names in The Index of Plant Names Index (IPNI) to the taxonomic literature.

## Materials and methods

The Index of Plant Names Index (IPNI, http://www.ipni.org) is an international register of published plant names based at the Royal Botanic Gardens, Kew but which has contributions from the Harvard Gray Index and the Australian Plant Name Index. Both new taxonomic names (e.g., for newly described species) and new combinations (e.g., reflecting transfers of species from one genus to another) are recorded in IPNI, together with a citation to the scientific work that published that name. These citations typically comprise an abbreviation for the publication (such as a a journal or a book), a description of the location of the name within that publication, such as a combination of volume number and page number, and the year of publication. One or more of these items may be missing, different journal abbreviations may be used in data sourced from different datasets, and the volume and pagination may be in either Roman or Arabic numerals. For some records the IPNI curators have added a link to the corresponding page in the Biodiversity Heritage Library (BHL), and for some recently added records the IPNI web site may give the DOI for a publication, but the majority of IPNI records are not linked to a digital identifier for the publication associated with each name.

In much the same way as for BioNames [Page 2013], I have harvested the IPNI database via its API and have developed software for matching the text string citations to digital identifiers such as DOIs, Handles, JSTOR links, etc. Whereas the source data for BioNames comprised citations at the level of a work (e.g., an article, chapter, or a book) which are relatively easy to match, the citations in IPNI are at the level of one or more pages within a work. Hence a big part of the challenge is to map page-level citations to work-level bibliographic data [Page 2009]. Given a complete bibliography of the taxonomic literature, this would be a relatively straightforward task, in that we could treat each work as comprising a set of pages, and we simply ask which works include the page in the IPNI citation. However, as yet we don’t have a comprehensive bibliography of life [King et al. 2011], hence much of the work in making the mapping involves scouring the web for sources of bibliographic information in the hope that these will include works containing the IPNI citations (the bibliographic database being assembled as a consequence of this work will be described in more detail elsewhere). I manage the mapping between IPNI names and the literature in a local MySQL database, and the results are periodically uploaded to a GitHub repository https://github.com/rdmpage/ipni-names, which also has code for a custom interface to that mapping.

## Datasette

A CSV file containing basic metadata for a plant name, such as IPNI LSID, scientific name, bibliographic details, and any identifiers found was generated from the current IPNI LSID to literature identifier mapping. This CSV file was then converted into a SQLite database using csv-to-sqlite, and the resulting database (ipni.db) was wrapped in a web server using the command “datasette serve ipni.db”. This datasets runs on the user’s local machine. We can also package the datasets into a Docker container using the following command:

~~~
datasette package -t <username>/ipni ipni.db
~~~

Where is your username at docker.com. The container can be run locally, or can be pushed to a repository where others can access it. To push to Docker’s Hub the commands are:

~~~
docker login -u <username> -p <password> docker push <username>/ipni
~~~

A container for this project is available at https://hub.docker.com/r/rdmpage/ipni/

## Results

The datasette generated here can be seen online at https://ipni.sloppy.zone. If this demo is offline, the reader can simply deploy a copy of the container from the Docker repository https://hub.docker.com/r/rdmpage/ipni/. The interface (Fig. 1) is simple and generic, but enables the user to browse the data as well as perform some straightforward queries. It should be noted that the interface can be customised to add more features, for this example I have stuck with the defaults.

**Figure 1:**
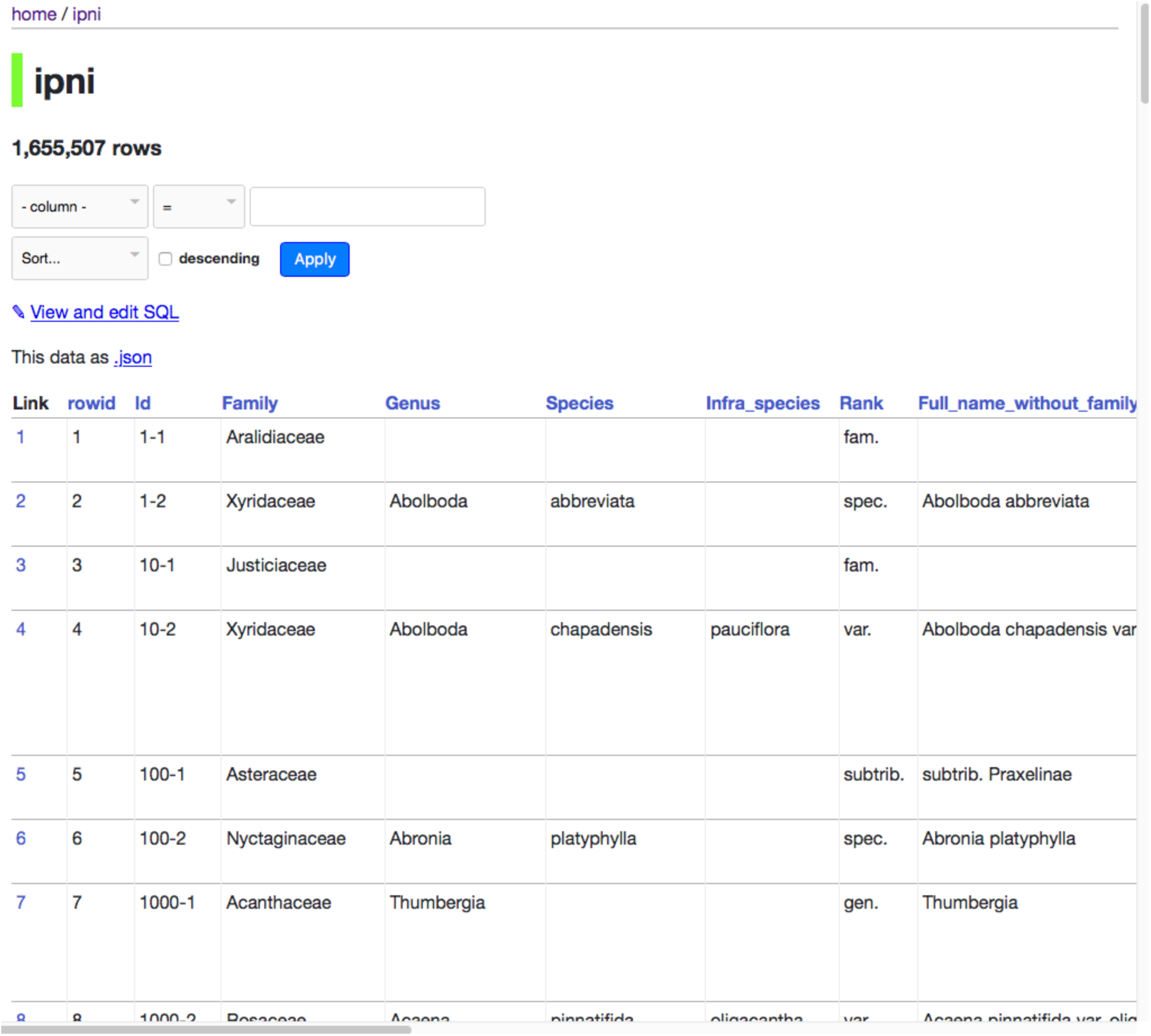
Screenshot of datasette of IPNI names.

Some simple queries include finding the DOIs for publications of new names in a given genus, such as *Begonia*:

~~~
select Id, Full_name_without_family_and_authors, doi from ipni where
~~~

~~~
Genus=“Begonia” and doi is not null;
~~~

JSTOR has digitised many botanical journals, so for some taxa such as the genus *Tiquilia* it is an excellent source of taxonomic literature:

~~~
select Id, Full_name_without_family_and_authors, doi from ipni where
~~~

~~~
genus=‘Tiquilia’ and jstor is not null;
~~~

Although the primary goal of the name-to-literature mapping is to find digital versions of the descriptions for each species, the datasette enables queries that might address other questions. For example, the database includes information on the agency that registers the DOI for a publication. For most publications this is CrossRef, but there are other agencies, such as DataCite, the multilingual European Registration Agency (mEDRA), and the Airiti Incorporation (華藝數位). Table 1 summarises the relative importance of these agencies. Different DOI agencies expose metadata for their DOIs in different ways, so the existence of multiple DOI agencies has implications for any researcher or tool that attempts to harvest bibliographic metadata. It could also can be used to investigate the pace of digitisation of legacy literature in different parts of the world. For example, a growing number of articles from journals published in China, Taiwan, and Japan now have DOIs assigned by local DOI agencies.

**Table 1:**
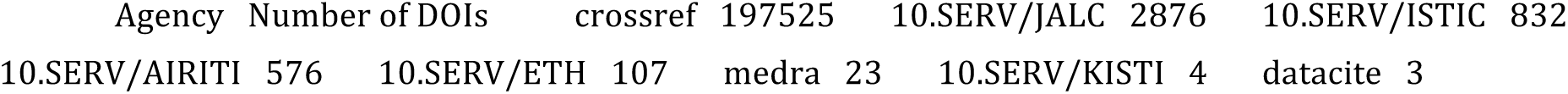
Number of DOIs for articles linked to IPNI names, grouped by registration agency.

## Discussion

Links between taxonomic names and the scientific literature have many possible uses. One is simply to be able to read the description of a new species, or discover the reasoning behind subsequent changes in name. Given that many of these sources are available in machine-readable text, the links could be used to generate a corpus for text mining to extract information on the species being described [e.g., Cui 2012].

The use of global bibliographic identifiers also enables queries that can span multiple databases. For example, knowing the DOI for a paper that changes the taxonomy of a plant genus, we could ask whether the evidence for that is supported by phylogenetic analysis by seeing whether that DOI also occurs in TreeBASE [https://treebase.org]. We could ask to what extent the discovery of new plants species is being driven by molecular data by seeing whether the DOI for the species description also occurs in sequence databases such as GenBank. However, these examples all require the existence of links between these databases, which are often incomplete [Miller et al. 2009], and hence represent further instances where the kind of mapping described here would be worthwhile.

In the absence of an existing knowledge graph, and the lack of a centralised infrastructure supporting its development, datasettes provide an easy mechanism for publishing links that places minimal burden on the researcher or curator doing the mapping, but also provides an interface that is potentially useful to users, even as we wait for the knowledge graph itself to coalesce.

